# Myosin II actively regulates *Drosophila* proprioceptors

**DOI:** 10.1101/2024.04.18.590050

**Authors:** Chonglin Guan, Kengo Nishi, Christian T. Kreis, Oliver Bäumchen, Martin C. Göpfert, Christoph F. Schmidt

## Abstract

Auditory receptors can be motile to actively amplify their mechanical input. Here we describe a novel and different type of motility that, residing in supporting cells, shapes physiological responses of mechanoreceptor cells. In *Drosophila* larvae, supporting cap cells transmit mechanical stimuli to proprioceptive chordotonal neurons. We found that the cap cells are strongly pre-stretched at rest to twice their relaxed length. The tension in these cells is modulated by non-muscle myosin-II motors. Activating the motors optogenetically causes contractions of the cap cells. Cap-cell-specific knockdown of the regulatory light chain of myosin-II alters mechanically evoked receptor neuron responses, converting them from phasic to more tonic, impairing sensory adaptation. Hence, two motile mechanisms seem to operate in concert in insect chordotonal organs, one in the sensory receptor neurons, based on dynein, and the other in supporting cells, based on myosin.

**One Sentence Summary:** Myosin II motors in contractile cells pre-stretch *Drosophila* stretch receptors.

## Main Text

Mechanoreceptors in animals detect physical forces from diverse origins such as touch, sound, fluid or air shear flow, ground vibrations or elastic deformations and convert these into electrical signals. Proprioception is a particular type of mechanosensation that depends on mechanosensory neurons distributed around muscles and joints that provide feedback about static and kinematic body positions. Proprioception is critical for coordinating muscle activity in locomotion (*1-3*). In mammals, stretch sensors encapsulating specialized intrafusal muscle fibers (muscle spindles) within skeletal muscles innervate primary sensory neurons (la afferents) and respond to muscle contractile status with bursts of action potentials. The proprioceptive neurons signal to the central nervous system (CNS) that then modifies movement commands to muscles through motor neurons (*4*). The signals sent to the CNS about muscle tension or joint position are dependent on how the cellular machinery of the sensors transmits mechanical inputs to mechanosensitive ion channels in the sensory neurons (*5-7*). Since channels are essentially digital sensors, either open or closed, mechanosensory machineries typically involve adaptation mechanisms that maintain an optimal working point even when mechanical boundary conditions change. In insects, proprioceptive chordotonal organs (ChOs), internal stretch receptor organs that are suspended under tension across joints or between body parts, sense conformational changes in the exoskeleton driven by muscle movements (*8, 9*). The most prominent ChOs of *Drosophila* larvae are the lateral pentascolopidial chordotonal organs (lch5) that are arranged serially along the larval abdomen (Fig. 1A) (*10*). Each lch5 organ is built of 5 parallel strands of 5 serially connected cells each, including a bipolar mechanosensory neuron that serves proprioception and vibration detection (*11, 12*). The organs are stretched between cuticle attachments via two sets of attachment cells, accessory ligament and cap cells (Fig. 1B). The ligament cells tie up the receptor somata, and the cap cells act as elastic cables transmitting force to the ciliated receptor dendrites tips (*13, 14*). The cap cells of lch5 organs are exceptionally long and contain actin cables and non-muscle myosin-II motors (*20, 24*), suggesting an active function.

**Fig. 1.**
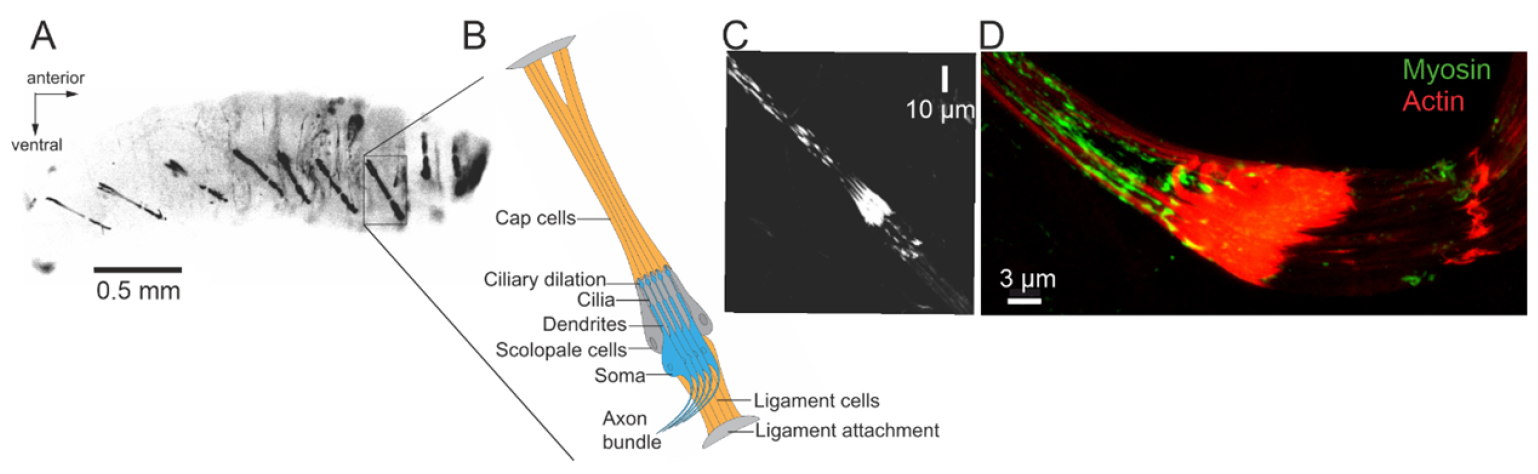
Anatomy of the larval pentascolopidial organ of *Drosophila*. **(A)** Strong GFP expression driven by *Tub85E-Gal4* in cap and ligament cells of lateral pentascolopidial organs (lch5) highlights their location on the ventral side of each larval hemisegment (contrast inverted) (see movie M1). **(B)** Schematic sketch of one lch5 organ, consisting of 5 parallel strings of cells, each comprising 6 cells. The bipolar chordotonal neurons (blue) are suspended between cap and ligament cells (orange) and anchored to the body wall by attachment cells (grey). The sensory dendrites are enveloped by scolopale cells (dark grey). **(C)** Phalloidin was used to label filamentous actin in the different cell types, making up an lch5 organ. (**D**) Myosin regulatory light chains were labeled by genetically attached GFP (*sqh-GFP*). GFP signals were enhanced with an anti-GFP antibody (green) and cells were counterstained for actin with phalloidin (red). Scale bars, (A) 0.5 mm; (B) 10 μm; (D) 3 μm.

The transient receptor potential (TRP) channels NOMPC, Nanchung, and Inactive, localized in the dendrites of lch5, have been implicated in mechano-electrical transduction (*15-19*). The lch5 organs are strongly pre-strained in their resting state (*20*), which is also evident from the straight, tense shape the organs maintain while the larva is moving (Movie M1). This construction makes it possible for the organ to detect both tension increases and decreases. To maintain optimal sensitivity in a moving animal, mechanosensory transduction typically involves adaptation mechanisms, which can involve active, force-generating processes. In *Drosophila* ChOs, the nature of adaptation mechanisms, however, had remained unknown. Myosins, a family of actin-binding motor proteins, are critical in a multitude of cellular processes that require forces and translocation (*21*). The mammalian myosin VII and myosin II have been linked to the sense of hearing and the function of inner-ear hair cells. Mutations in these motors can cause severe neurosensory pathologies such as Usher syndrome (*22*). In the fly hearing system, studies have implicated a crucial role of actomyosin-based structures in the architecture of ChOs (*23, 24*). The actual molecular mechanisms of myosin-dependent tension regulation in mechanosensing, however, remained elusive, because the functional dissection of ChO mechanics *in vivo* has been challenging. We have mechanically measured lch5 organ elasticity and have probed for contributions of myosin activity. We furthermore electrically recorded from the axons of the sensory neurons during mechanical stimulation and again probed for the role of myosin.

To quantify lch5 organ elasticity, we used a filet prep of third instar larvae (*25*), mounted under buffer solution in an upright microscope (Fig. 2B). To stretch an organ, we laterally displaced its midpoint with a calibrated micropipette force sensor (*26*). This procedure stretched the organ into a triangular shape (Fig. 2A), different from the organ’s axial stretch during larva movement, where cuticle deformation changes the distance between the two attachment points of the organ (Movie M2). Since, however, bending is localized to the point of contact between micropipette and organ, we assume that the mechanical actuation of the sensory dendrite is similar to the one it experiences in the moving animal. Note that the lateral stretching experiment simultaneously measures the pretension in the organ and the elastic modulus (Eqs. 4 and 5). The primary data resulting from the experiment is the micropipette displacement normal to the initial orientation of the organ at a given micropipette force. The displacement increased almost linearly with increasing force, yielding an effective spring constant of 31.7 ± 1.9 mN/m for lch5 organs with lengths between 394 and 424 μm in wildtype larvae (Fig. 2D). We found an effective Young’s modulus of the organ of ∼300 kPa and a strong ∼100% prestrain in third instar larvae. The neuronal response to stretch showed rapid adaptation. Cap-cell-specific knockdown of the regulatory light chain of myosin-II led to more tonic neural responses, reflecting alterations in mechanosensory adaptation.

**Fig. 2.**
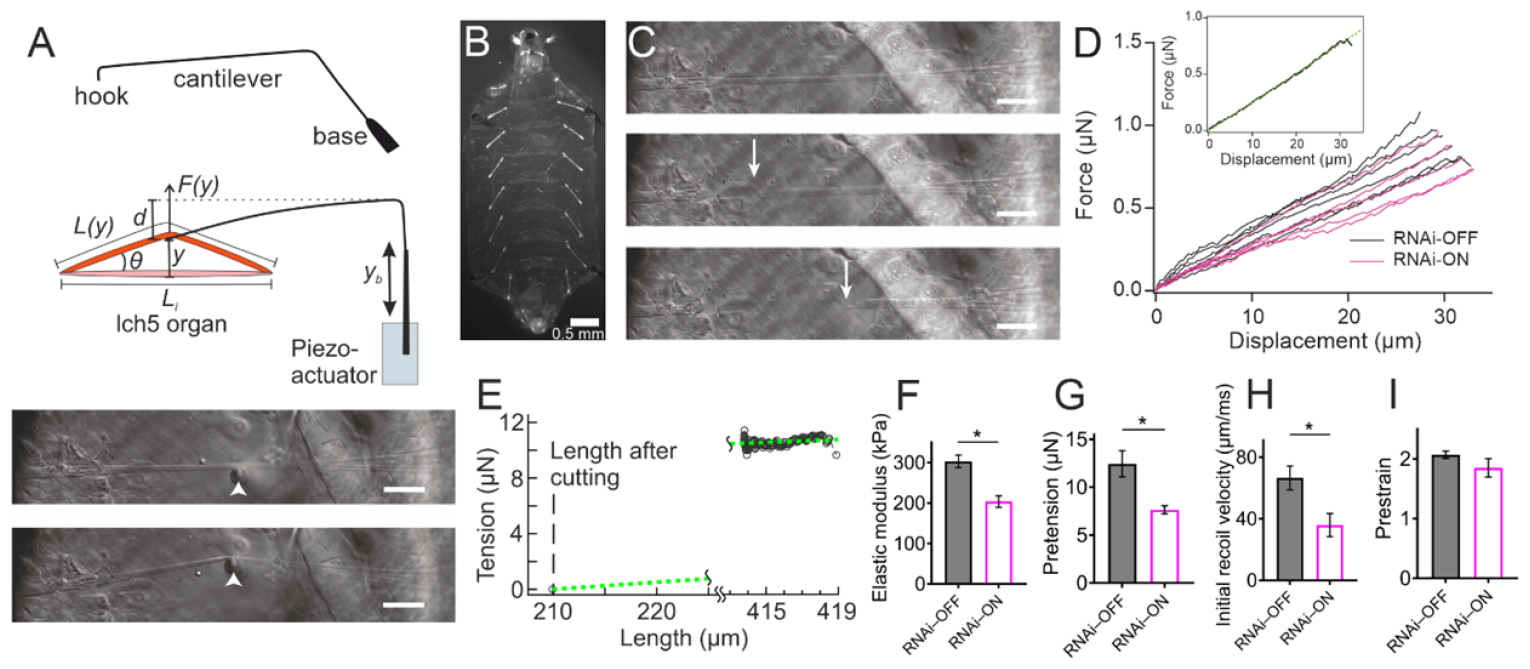
Elasticity, pre-strain, and pretension of lch5 organs and the effect of myosin inactivation. **(A)** Schematic of the glass micropipette micromanipulation experiment. Top: The perspective view shows the glass pipette twice bent by 90° with the thin long cantilever segment acting as a calibrated spring. Middle: Top view: axial movement of the base of the micropipette by a distance of *y*_*b*_ results in bending of the cantilever and a deflection of the tip of the micropipette and the midpoint of the organ by a distance of *y*. The initial organ length *L*_*i*_ is extended to the stretched length *L*(*y*). The bend amplitude of the cantilever is calculated as *d = y*_*b*_ *– y*, from which the force on the organ is calculated. Bottom: lateral deflection of the lch5 organ (white arrows mark the tip of the micropipette). Scale bar: 50 μm. **(B)** Filet preparation for probing lch5 organ mechanics. GFP expression driven by *Tub85E-Gal4* in cap and ligament cells of lch5 shows their symmetrical localization on the ventral side of each larval hemisegment. **(C)** DIC images of UV-laser ablation experiments. Shown here are (top) before ablation, (middle) 1 frame (1 ms) after UV-laser pulse, and (bottom) relaxed state. Arrows show the end of lch5 organ fragment during recoiling. Scale bar: 50 μm. **(D)** Force-displacement curves of control organs (RNAi-OFF) and myosin-inactivated organs (RNAi-ON), deflected normal to their long axis near their midpoints (N = 6 larvae for control organs and N = 5 larvae for myosin-inactivated organs). RNAi-OFF and RNAi-ON are shown as the black and magenta lines. Inset: a typical force curve of RNAi-OFF is plotted with the fitting results using Eq. 4 (dotted green line). **(E)** A representative tension curve of a control animal is plotted with the fitting results using Eq. 5 (dotted green line), using *G* obtained from the previous fitting analysis with Eq. 4 and *L*_0_ from the UV cutting experiment, respectively. Extrapolation of organ length to zero tension results in a resting length consistent with the results of laser cutting experiments (∼210 μm). **(F-I)** Comparison of parameters from controls and myosin-inactivated lch5 organs. **(F)** Young’s moduli; **(G)** Pre-tension; **(H)** Initial recoil velocity; **(I)** Pre-stain. Data are presented as mean ± SEM. Asterisks denote comparisons of values with a two-tailed Mann-Whitney test, *p ≤ 0.05.

In order to calculate the initial pretension and the (coarse-grained) elastic modulus of the organ from the force-displacement curves, we assumed that lch5 can be described as a pre-strained bundle of 5 strings, each with a homogeneous elastic Young’s modulus *E*. With this assumption, we average over the internal material complexity of the 5 types of cells the lch5 organs are built from, as well as over their mechanical differences. The longest (∼ 70% of the total organ length) and most stretchable cells in lch5 organs are the cap cells. To describe the pre-strain, we assume that the initial resting length *L*_*i*_ is longer than the organ’s relaxed length *L*_*0*_. When the midpoint of the lch5 organ is laterally moved by a distance *y* due to the lateral force *F*(*y*), the elastic energy *U*(*y*) stored in the organ can be written as:

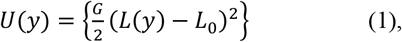

with *L*(*y*) being the contour length of the organ at displacement *y. G* is an elastic coefficient that can be related to the coarse-grained Young’s modulus *E* by *G* = 5*Eπa*^2^/*L*_0_. We here assume that each string of cells can be approximated as a cylinder with radius *a*. The factor 5 arises because lch5 comprises five parallel strings of cells that each include one sensory neuron associated with one supporting cap cell. Since the organ is pre-strained, the initial conformation is straight as are the two legs of the triangle after stretching (Fig. 2A Middle, Bottom). We can therefore obtain a geometric constraint:

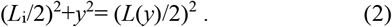

Under this condition, we find the force the organ exerts on the needle when its midpoint is laterally displaced by *y* by taking a derivative of the energy *U* with respect to *y*:

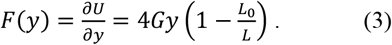

By combining Eqs. 2 and 3, the force can be expressed as:

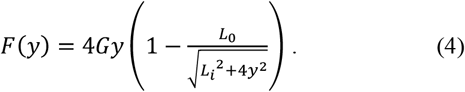

For small *y*, Eq. 4 can be approximated as 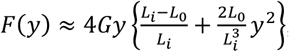, implying that the force initially increases approximately linearly with *y* in case the organ is pre-strained, *L*_0_ > *L*_*i*_. This linear term vanishes without pre-strain, i.e. when *L*_0_ = *L*_*i*_. Furthermore, we can estimate the axial tension in the organ resulting from the lateral deflection. Defining an angle *θ* between the legs of the triangle and the line connecting the organ’s attachment points to the cuticle (Fig. 2A Middle), the tension *T* is calculated as:

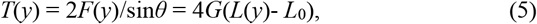

where *θ* fulfills sin*θ* = 2*y*/*L* and we used Eq. 3 in the last step. Pretension is defined as the tension before stretching, i.e., the tension at *y* = 0 that we can obtain by extrapolation to y = 0. We fitted force-displacement curves of lch5 organs using Eq. 4 to estimate their elastic modulus. To reduce the number of fit parameters, we determined *L*_0_ of each organ by UV-laser cutting after performing the stretching experiments (Movie M3). As shown in Fig. 2C, lch5 organs with initial lengths between 394 μm and 424 μm rapidly retracted after UV cutting by distances between 194 μm and 243 μm, resulting in an average shrinkage of 51.5% ± 1.4%. The initial recoil velocity was 67 ± 8 μm/ms (Fig. 2H). This result quantifies the substantial pre-strain in the organs, *L*_i_/*L*_0_ ∼ 2. Eq. 4 closely fits the experimental force-displacement curve (Fig. 2D), supporting the validity of our model. Fitting force-displacement curves from 6 control larvae yielded an elastic Young’s modulus of *E* = 303 ± 15 kPa (Fig. 2F), which is higher than that of muscle (∼ 1 kPa) but lower than that of tendon (∼10 MPa) (*27-29*). Given the measured pre-strain, this elastic modulus means that the pretension is 12.4 ± 1.4 μN (Fig. 2G). Recently, Hassan *et al*. reported an elastic modulus of cap cells of 9.5 ± 0.3 kPa from AFM measurements (*30*), a value 30 times smaller than our result. It should be noted that an AFM cantilever locally indents the surface of organs with difficult to determine mechanical boundary conditions, while our micropipette experiment globally stretches the whole organ, which comes closer to natural conditions and likely explains the difference. As shown in Fig. 2E, we found that tension remains relatively constant, as effectively reproduced by Eq. 5, within the range of displacements applied via micropipettes. This is due to the fact that the relative length changes of the organ induced by the micropipette force sensors are much smaller than its pre-strain. It is interesting to note that this construction principle, using a strongly prestrained elastic element to monitor small additional length changes, produces a sensor that works at almost constant tension. The almost constant tension facilitates adaptation mechanisms that maintain maximal sensitivity (channel half-maximum open probability) of lch5 organs during larval crawling.

The cap cells of lch5 organs contain actin bundles and non-muscle myosin-II motors in arrangements resembling stress fibers (Fig. 1C and D), suggesting a possible myosin-dependent contractile function. A system generating a variable tension in the lch5 organs would be a prime candidate for an adaptation mechanism. We thus set out to measure sensory adaptation. A first indication that adaptation takes place was seen in experiments where an organ was rapidly deflected laterally in a stepwise fashion. We always observed a slow relaxation of the tension after such deflections (Fig. 3C). To assess the effect of myosin-II on lch5 mechanics, we conducted a knockdown experiment targeting Spaghetti Squash (SQH), the regulatory light chain of non-muscle myosin-II, which is highly expressed in cap cells (*20*). SQH was down-regulated using RNA interference, accomplished by driving UAS-*sqh*-RNAi (*32*) with *pinta*-GAL4 (*33, 34*). UV-laser cutting experiments showed that the knockdown of myosin II significantly reduced retraction amplitude and recoil velocity (36 ± 8 μm/ms (Fig. 2H)) at a pre-strain of 1.85 ± 0.16 (Fig. 2I). Fitting lateral stretching experiments with Eq. 4 resulted in a Young’s modulus of myosin II knockdown organs of 204 ± 14 kPa (Fig. 2F), and a pre-tension of 7.6 ± 0.4 μN (Fig. 2G). Both values are significantly smaller than those observed in control larvae. These results demonstrate that the myosin activity in cap cells, interacting with actin filaments, affects both the material properties and the mechanical tension of lch5 organs.

**Fig. 3.**
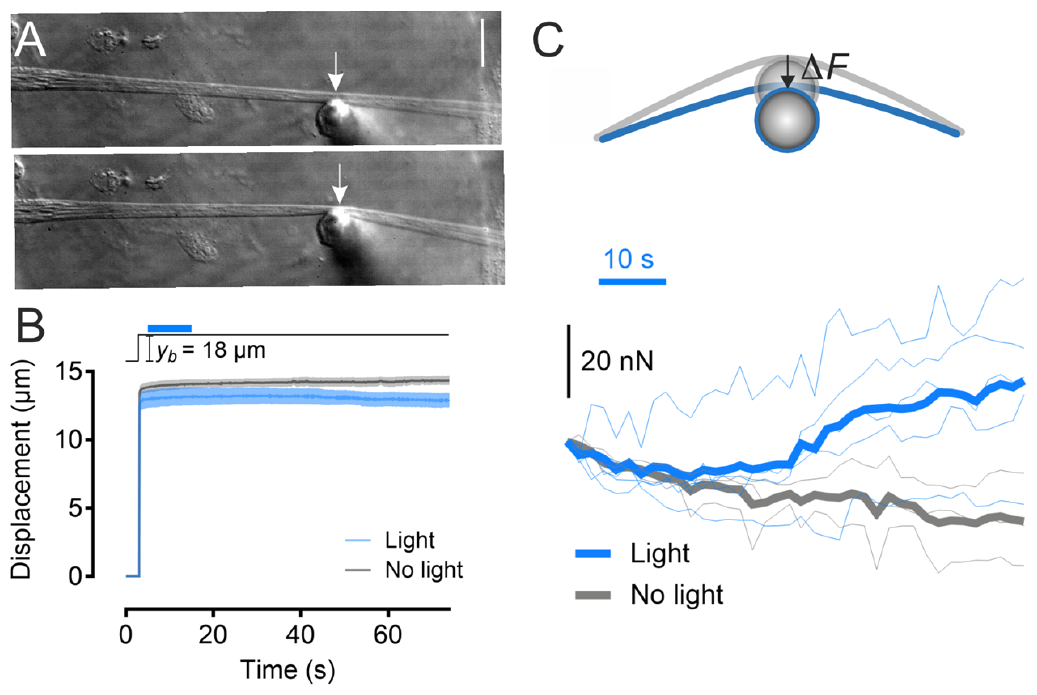
Optogenetic activation of Myosin II promotes lch5 contractions. **(A)** DIC images of lch5 organ stretched by ∼10 μm deflection of its midpoint with a micropipette (white arrows). Top: undeflected, bottom: deflected. Force is calculated from the position of the pipette tip. Scale bar: 20 μm. **(B)** Top: Displacement of the base of the micropipette (black line) and 10 s illumination period (blue bar). Bottom: Displacement of the pipette tip over time with (blue/light blue, average ± SEM, N = 5 larvae) and without (grey/black, average ± SEM, N = 3 larvae) light-induced optogenetic myosin-II activation. **(C)** Top: Schematic of contraction of the organ leading to micropipette tip movement. Bottom: Force as a function of time with (blue: individual curves; dark blue: average) and without (grey: individual curves; dark grey: averages) light-induced optogenetic myosin-II activation. Light was on continuously for 10 s (blue bar).

To confirm the role of myosin in generating tension in the cap cells, we employed an optogenetic strategy. Phosphorylation of its regulatory light chain activates myosin-II motors. This phosphorylation can be induced optogenetically by light-activating Rho1 signaling using the RhoGEF2-CRY2/CIBN system (*35, 36*). We expressed this system in lch5 cap cells with the driver *pinta-Gal4* to optogenetically activate myosin-II motors. In order to quantify the effect, we laterally deflected lch5 at its midpoint by ∼10 μm with a calibrated glass micropipette with and without illumination. By monitoring the position of the pipette tip, we determined the force exerted by the organ on the pipette (Fig. 3A). In the absence of light, this force slowly declined by on average 0.25 nN per second (Fig. 3C grey trace), reflecting slow chordotonal organ relaxation. When we light-activated myosin-II in the cap cells, however, the force increased with an average slope of 0.35 nN per second for tens of seconds, demonstrating that the activity of myosin-II motors triggers cap cell contractions (Fig. 3B, C; Fig. S1).

Given that myosin-II motors promote contractions of cap cells and contribute to organ pretension and elasticity, we wondered how the motors contribute to the mechanosensory organ function. lch5 comprises five mechanoreceptor cells that spike spontaneously and increase their spiking frequency upon mechanical stimulation (*25, 37*). To examine whether myosin activity in cap cells affect the electrical responses of the neurons, we recorded the neuron activities extracellularly by sucking the axon bundle into a recording electrode (Fig. 4A) while light-activating myosin II in the cap cells. We found that the neuron firing rate increased approximately 25% during the light application and returned to baseline after illumination (Fig. S1B). Hence, the mechanoreceptors are sensitive to myosin-driven lch5 contractions, providing a potential adaptation mechanism. We further tested this hypothesis by knocking down *sqh* in the cap cells and measuring mechanically evoked receptor responses. We again laterally deflected lch5 at its midpoint with a rigid glass needle while recording action currents from the lch5 receptor neurons (Fig.4A; Fig. S2A, D). The tip of the glass needle, driven by a piezo-actuator, was placed on one side of the organ and was used to apply transient mechanical loads. Stepping up and down the tension in the organs evoked predominantly phasic responses in RNAi-OFF controls, with spiking frequencies increasing transiently after the start and end of the deflections. Knocking down *sqh* in the cap cells converted these predominantly phasic spiking responses into tonic ones, with the spiking frequency remaining increased during the entire stimulus presentation (Fig. 4B, C). A slowing down of current transients, as seen in RNAi-ON larvae, might reflect alterations in mechanoreceptor adaptation. To isolate the underlying receptor currents, we blocked voltage-gated sodium channels with tetrodotoxin (TTX) to prevent action current firing. In RNAi-OFF controls, stimulus start and end evoked transient current responses that declined gradually, but not completely, to the steady state level seen before the stimulus, consistent with the predominantly phasic action potential firing (Fig. S2B, D). Fitting the transient current responses with an exponential yielded an adaptation time constant of 44 ms. In RNAi-ON larvae, the decline of the current response after start and end of a stimulus was slowed down, resulting in a substantially larger time constant of 102 ms (Fig. 4D, H). Myosin-II thus modulates both lch5 mechanics and lch5 mechanosensory responses (Fig. 4E, F, G and H). We further probed adaptation by imposing a series of force steps onto the organ before and after deflecting it with offset displacements, allowing 2 s for adaptation. Compared to the current-displacement (I/X) relation before the offset displacement, those measured after application of offset displacement were shifted to larger displacements (Fig. S3A, B and C). Plotting this adaptive shift against the offset displacement revealed that, in both RNAi-On and –Off larvae, lch5 mechanoreceptors adapt incompletely. The degree of adaptation was lowered in RNAi-On larvae (75% compared to RNAi-OFF controls (84%) (Fig. 4I). Hence, myosin-II in lch5 cap cells contributes to mechanosensory adaptation.

**Fig. 4.**
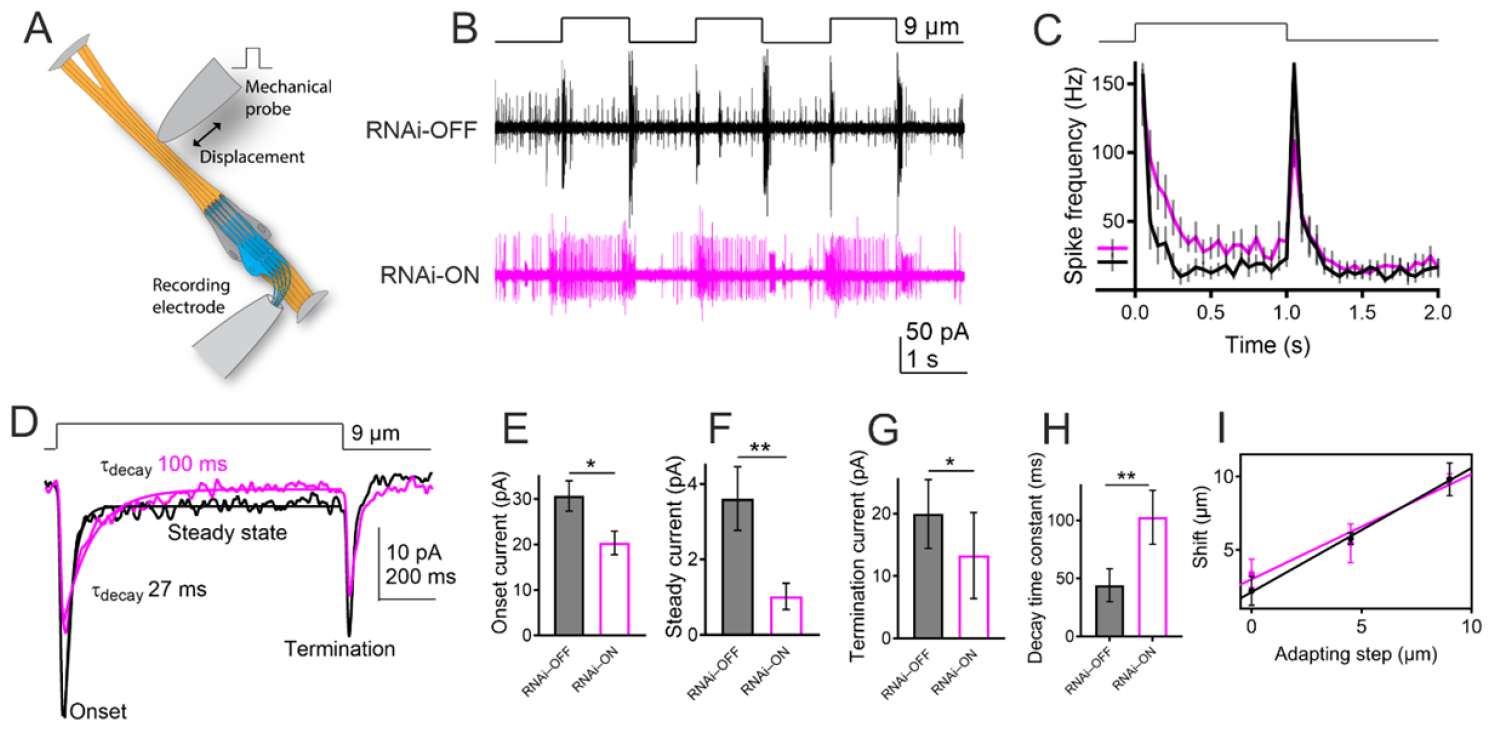
Mechano-electrical transduction of lch5 neurons. **(A)** Schematic of experiment of simultaneous mechanical stimulation of lch5 organs and electrical recording from the axons of the sensory neurons. The organ was put under increased tension by transverse deflection of the cap cell bundle with a rigid glass needle mounted on a piezo actuator. Severed axons of all 5 sensory neurons were sucked into a recording pipette electrode connected to a patch-clamp amplifier. **(B)** Spiking responses in control (RNAi-OFF: black) and sqh-knockdown (RNAi-ON: purple) animals to 9 μm displacements, generating about 12 nN tension increase. **(C)** Action current frequencies (50-ms bins) after start and end of mechanical stimulation. (RNAi-OFF: black, RNAi-ON: purple). **(D)** Averaged receptor currents of control (N = 10) and sqh knockdown (N = 9). RNAi-ON larvae displayed significantly lower peak responses to onset, steady state and termination of mechanical loads. **(E)**, steady-state **(F)** and termination **(G)** of mechanical loads. **(H)** The time constant of adaptation for RNAi-ON organs was significantly increased. **(I)** The extent of the adaptive response was impaired by myosin inactivation (slope = 0.72) in comparison with that of control organs (slope = 0.84). Data are presented as mean ± SEM. Asterisks denote comparisons of values with a two-tailed Mann-Whitney test, *p ≤ 0.05, **p ≤ 0.001.

In conclusion, we have shown *in vivo* that myosin motors in the supporting cap cells affect both mechanical properties and mechanically evoked responses of *Drosophila* lch5 chordotonal organs. We discovered a crucial role of motors in sensory adaptation and could show how tension-generating mechanisms contribute to the physiology of mechanoreceptor neurons. We found lch5 organs to be extremely elastic, ultimately supporting more than 200% strain. A 100% pre-strain in third instar larvae keeps tension relatively stable while the animal is moving, presumably aiding sensory adaptation. Adaptation occurs on the scale of tens of ms, creating a high-pass filter characteristic for the proprioceptive response of the organs, presumably filtering out slow shape changes when the larva is crawling. We found that the myosin-dependent adaptation mechanism resides in the supporting cap cells, where actomyosin structures resembling stress fibers can contract, affecting the neuronal response. A stress sensing machinery typically includes a transducer channel that is activated by force, and active (visco-)elastic elements that serve to adapt the sensor to changing mechanical boundary conditions and thereby maintain maximal sensitivity. We hypothesize a simple mechanical model for lch5 organs (Fig. S4): The extracellular matrix around the organ acts as an elastic spring providing the largest part of the observed pre-stress. A second adaptation spring composed of active, force-generating non-muscle myosin motors and filamentous actin structures in cap cells regulates resting tension and the gating forces acting on mechanosensitive channels in the sensory dendrites. Given that dynein-driven cilium motility boosts mechanosensitivity by amplifying vibrations (*38-40*), our results indicate that in insect chordotonal organs, two active processes cooperate: dynein activity and myosin-based cap cell contractility. More work is needed in the future to address communication between the two processes. Larval lch5 organs provide a promising opportunity to study the molecular mechanisms involved in the regulation and encoding of mechanical forces by primary mechanoreceptor neurons and to explore the roles of various mechanical components, such as extracellular matrix, molecular motors, and the cytoskeleton, in controlling cell mechanics and sensory transduction.

## Acknowledgments

This work was supported by grants from the European Research Council (ERC) under the European Union’s Seventh Framework Programme (FP7/2007-2013) (grant agreement n°340528) (CS), and from the HFSP (RGP009/2023) (M.C.G.), and from the Multiscale Bioimaging Cluster of Excellence (MBExC), University of Göttingen (M.C.G.) and by the Max Planck Society (OB). Additional data, including raw data, are presented in the supplementary materials. The authors thank Daniel Kiehart and Jörg Albert for valuable discussions.

## Author contributions

Conceptualization: CG, MCG, CFS

Methodology: CG, KN, CTK, OB

Investigation: CG, KN

Visualization: CG, KN, CTK, OB, MCG

Funding acquisition: CFS, MCG

Supervision: MCG, CFS

Writing – original draft: CG, KN

Writing – review & editing: MCG, CFS, OB

## Supplementary Materials

Materials and Methods

Supplementary Text

Figs. S1 to S4

References (*41-44*)

Movies M1 to M3

## References and Notes

1. S. H. Scott, Optimal feedback control and the neural basis of volitional motor control. Nat. Rev, Neurosci. 5, 532–546 (2004).

2. U. Windhorst, Muscle proprioceptive feedback and spinal networks. Brain Res. Bull. 73, 155–202 (2007).

3. J. C. Tuthill, E. Azim, Proprioception. Curr. Biol. 28, R194–R203 (2018).

4. A. Crowe, P. B. Matthews, The effects of stimulation of static and dynamic fusimotor fibres on the response to stretching of the primary endings of muscle spindles. J. Physiol. 174, 109–131 (1964).

5. M. Chalfie, Neurosensory mechanotransduction. Nat. Rev Mol. Cell Biol. 10, 44–52 (2009).

6. C. C. DuFort, M. J. Paszek, V. M. Weaver, Balancing forces: architectural control of mechanotransduction. Nat. Rev. Mol. Cell Biol. 12, 308–319 (2011).

7. D. E. Ingber, N. Wang, D. Stamenovic, Tensegrity, cellular biophysics, and the mechanics of living systems. Rep. Prog. Phys. 77, 046603 (2014).

8. L. H. Field, T. Matheson, Chordotonal organs of insects. Adv. Insect Physiol. 27, 1–228 (1998).

9. J. C. Tuthill, R. I. Wilson, Mechanosensation and adaptive motor control in insects. Curr. Biol. 26, R1022–R1038 (2016).

10. V. Orgogozo, W. B. Grueber, FlyPNS, a database of the Drosophila embryonic and larval peripheral nervous system. BMC Dev. Biol. 5, 4 (2005).

11. J. C. Caldwell, M. M. Miller, S. Wing, D. R. Soll, D. F. Eberl, Dynamic analysis of larval locomotion in Drosophila chordotonal organ mutants. Proc. Natl. Acad. Sci. U.S.A. 100, 16053–16058 (2003).

12. W. Zhang, Z. Yan, L. Y. Jan, Y. N. Jan, Sound response mediated by the TRP channels NOMPC, NANCHUNG, and INACTIVE in chordotonal organs of Drosophila larvae. Proc. Natl. Acad. Sci. U.S.A. 110, 13612–13617 (2013).

13. M. J. Kernan, Mechanotransduction and auditory transduction in Drosophila. Pflugers Arch. 454, 703–720 (2007).

14. M. C. Göpfert, R. M. Hennig, Hearing in Insects. Annu. Rev. Entomol. Vol 61 61, 257–276 (2016).

15. T. Effertz, R. Wiek, M. C. Göpfert, NompC TRP channel is essential for Drosophila sound receptor function. Curr. Biol. 21, 592–597 (2011).

16. D. Zanini, M. C. Göpfert, TRPs in hearing. Handb. Exp. Pharmacol. 223, 899–916 (2014).

17. M. C. Göpfert, J. T. Albert, B. Nadrowski, A. Kamikouchi, Specification of auditory sensitivity by Drosophila TRP channels. Nat. Neurosci. 9, 999–1000 (2006).

18. T. Effertz, B. Nadrowski, D. Piepenbrock, J. T. Albert, M. C. Göpfert, Direct gating and mechanical integrity of Drosophila auditory transducers require TRPN1. Nat. Neurosci. 15, 1198–1200 (2012).

19. B. P. Lehnert, A. E. Baker, Q. Gaudry, A. S. Chiang, R. I. Wilson, Distinct roles of TRP channels in auditory transduction and amplification in Drosophila. Neuron 77, 115–128 (2013).

20. A. Prahlad et al., Mechanical properties of a Drosophila larval chordotonal organ. Biophys. J. 113, 2796–2804 (2017).

21. M. Vicente-Manzanares, X. Ma, R. S. Adelstein, A. R. Horwitz, Non-muscle myosin II takes centre stage in cell adhesion and migration. Nat. Rev. Mol. Cell Biol. 10, 778–790 (2009).

22. C. Bonnet, A. El-Amraoui, Usher syndrome (sensorineural deafness and retinitis pigmentosa): pathogenesis, molecular diagnosis and therapeutic approaches. Curr. Opin. Neurol.25, 42–49 (2012).

23. S. V. Todi, J. D. Franke, D. P. Kiehart, D. F. Eberl, Myosin VIIA defects, which underlie the Usher 1B syndrome in humans, lead to deafness in Drosophila. Curr. Biol. 15, 862–868 (2005).

24. T. Li et al., The E3 ligase Ubr3 regulates Usher syndrome and MYH9 disorder proteins in the auditory organs of Drosophila and mammals. eLife 5, (2016).

25. N. Scholz et al., The adhesion GPCR latrophilin/CIRL shapes mechanosensation. Cell Rep. 11, 866–874 (2015).

26. C. T. Kreis, M. Le Blay, C. Linne, M. M. Makowski, O. Baumchen, Adhesion of Chlamydomonas microalgae to surfaces is switchable by light. Nat. Phys. 14, 45–49 (2018).

27. K. Arda, N. Ciledag, E. Aktas, B. K. Aribas, K. Kose, Quantitative assessment of normal soft-tissue elasticity using shear-wave ultrasound elastography. AJR Am. J. Roentgenol. 197, 532–536 (2011).

28. B. C. W. Kot, Z. J. Zhang, A. W. C. Lee, V. Y. F. Leung, S. N. Fu, Elastic modulus of muscle and tendon with shear wave ultrasound elastography: variations with different technical settings. PloS One 7, (2012).

29. Y. N. Feng, Y. P. Li, C. L. Liu, Z. J. Zhang, Assessing the elastic properties of skeletal muscle and tendon using shearwave ultrasound elastography and MyotonPRO. Sci. Rep. 8, 17064 (2018).

30. A. Hassan et al., A change in ECM composition affects sensory organ mechanics and function. Cell Rep. 27, 2272–2280 e2274 (2019).

31. P. Jordan, R. Karess, Myosin light chain-activating phosphorylation sites are required for oogenesis in Drosophila. J. Cell Biol. 139, 1805–1819 (1997).

32. L. A. Perkins et al., The transgenic RNAi project at Harvard Medical School: resources and validation. Genetics 201, 843–852 (2015).

33. R. Katana et al., Chromophore-independent roles of opsin apoproteins in Drosophila mechanoreceptors. Curr. Biol.: CB, (2019).

34. D. Zanini et al., Proprioceptive opsin functions in Drosophila larval locomotion. Neuron 98(1):67-74.e4. (2018).

35. E. Izquierdo, T. Quinkler, S. De Renzis, Guided morphogenesis through optogenetic activation of Rho signalling during early Drosophila embryogenesis. Nat. Commun. 9, 2366 (2018).

36. J. R. Sellers, Regulation of cytoplasmic and smooth muscle myosin. Curr. Opin. Cell Biol. 3, 98–104 (1991).

37. N. Scholz et al., Mechano-dependent signaling by Latrophilin/CIRL quenches cAMP in proprioceptive neurons. eLife 6:e28360. (2017).

38. M. C. Göpfert, D. Robert, Motion generation by Drosophila mechanosensory neurons. Proc. Natl. Acad. Sci. U.S.A. 100, 5514–5519 (2003).

39. B. Nadrowski, J. T. Albert, M. C. Göpfert, Transducer-based force generation explains active process in Drosophila hearing. Curr. Biol. 18, 1365–1372 (2008).

40. S. Karak et al., Diverse roles of axonemal dyneins in Drosophila auditory neuron function and mechanical amplification in hearing. Sci. Rep. 5, 17085 (2015).

