## Supplementary for "Myosin II actively regulates *Drosophila* proprioceptors"

### Materials and Methods

#### Fly stocks

Third instar larvae were raised on standard cornmeal medium at 25°C and 75% humidity. The following strains were used in this study:

*$w^{1118}$* (Bloomington Drosophila Stock Center, # BL3605)

*pinta-GAL4* (Göpfert lab)

*$w^{1118}$; P{sqh-GFP.RLC}3* (Bloomington Drosophila Stock Center, BDSC#57145)

*UAS-YFP; Tub85E-Trojan-GAL4/sb* (Göpfert lab)

*UAS-sqhRNAi/Cyo* (Bloomington Drosophila Stock Center, BDSC#38222)

*If/CyO; UAS-RhoGEF2-CRY2::mCherry]/TM3, Ser and UAS-CIBN::pmGFP]/Cyo; Sb/TM3, Ser.* (De Renzis lab)

#### Preparation of larvae/fillets

A third-instar larva was pinned on a PDMS block in a Petri dish. A larval fillet was cut in hemolymph-like saline (103 mM NaCl, 3 mM KCl, 4 mM $MgCl_2$, 5 mM TES, 7 mM sucrose, 10 mM glucose, 10 mM trehalose, 26 mM $NaHCO_3$, 1 mM $NaH_2PO_4$, 2 mM $CaCl_2$, pH to 7.25). Hemi-segments A3–A6 of the larval fillet were selected for further dissection. The major muscles covering the body wall were crosscut along the center of their length. Extra muscles were removed gently with fine scissors to expose the ChO organs. The fillet preparation was then transferred to a microscope stage for electrophysiological/mechanical measurements.

### Micropipette force spectroscopy

Micropipette force spectroscopy (MFS) is a technique based on the deflection of a thin glass needle (micropipette) that was successfully applied in the past to measure forces and mechanical properties in various biological systems *(43)* such as rupture forces of actin filaments (*41*), adhesion forces of unicellular microalgae (*44*), and elastic properties of the nematode *C. elegans* (*42, 43*). The force measurement is based on the high-resolution optical imaging of the deflection of a custom-shaped and calibrated micropipette force sensor.

As force sensors, we used micropipettes fabricated from borosilicate glass capillaries (WPI, Borosilicate Glass Capillaries TW100-6, initial outer diameter 1 mm) as described earlier (*43*, *44*). A P-97 Flaming/Brown Micropipette Puller (Sutter Instrument) was used to pull pipettes. These straight micropipettes of several centimeter length and outer diameter of 20-30 µm were then bent into the final shape depicted in Figure 2A using a microforge (Narishige, Microforge MF-900). The resulting cantilevers had a length of approximately 1 to 2 cm, which was at least one to two orders of magnitude longer than the short vertical hook that was used to contact an lch5 organ. This design guaranteed that the short hook was much stiffer than the cantilever arm, such that any deflection of the hook could be neglected. Furthermore, no substantial torque within the cantilever arm was observed during any experiment, since the tip of the micropipette remained in focus during measurement, indicating that the z-position of the tip did not change. The position of the tip was tracked from recorded movies by the Template Matching plugin in ImageJ.

### Force-sensor calibration

The micropipette force sensors were calibrated using a second pre-calibrated micropipette force sensor. This reference cantilever featured a double-L-shape and was calibrated using the added weight of a variable mass, as described earlier (*43*). The spring constant was obtained from the mean of multiple independent calibration experiments. The spring constants of the force sensors employed in this study varied from 10 to 70 nN/µm. Independent validations of the micropipette calibration with the added weight method confirmed the spring constants obtained by the reference cantilever method.

### Laser ablation

UV-laser ablations were performed using a pulsed diode-pumped solid-state laser (355 nm, repetition rate 1 kHz, pulse energy ~42 µJ, pulse length <1.5 ns) attached to an upright microscope (Zeiss Axio Examiner Z) using a 20X water-immersion objective (NA = 1). To perform ablations, the laser was targeted to the boundary line between scolopale cells and cap cells. Differential interference contrast images were collected by a high-speed camera (FASTCAM SA1.1, Photron) every 1 ms.

### Electrophysiological recordings

During preparation of filets, larvae were immersed in hemolymph-like saline. Muscles covering chordotonal neurons were gently removed with fine scissors. The axon bundle was severed using a UV-laser (DPSL-355/42, UGA-42 Caliburn Ablation System, Rapp Optoelectronic). The activity of lch5 neurons was recorded from the axon bundle using a suction electrode (GB1508P, Science Products) pulled and fire polished with a DMZ Universal Puller (Zeitz Instruments) to a tip diameter of approximately 5 µm. An extracellular amplifier (EXT-02B, npi electronic GmbH) and a patch-clamp amplifier (ELC-03XS, npi electronic GmbH) were coupled to a digitizer (LIH 8+8, HEKA Elektronik) for event frequencies and receptor-current recordings, respectively.

Mechanical stimulation was applied through a piezo-actuated (P-841.3 Actuator and E-709 Controller, Physik Instrumente GmbH) custom-made glass needle, placed approximately 100 µm away from the scolopale cells on a side of the cap cell bundle. Cap cells were pushed laterally with various step displacements. Spontaneously active neurons were stimulated either optogenetically or mechanically using the indicated step displacements (three cycles of 1 s stimulation preceded by 1 s rest for each step). Data were filtered with a Bessel filter at 2.9 kHz and sampled at 10 kHz. The dynamics of event frequencies was analyzed using custom-made MATLAB (Math Works) programs. To access receptor currents, 4 µm TTX was added to the bath solution to prevent neuronal firing. Current traces were analyzed in Clampfit 10.2 (Molecular Devices). Before measuring the amplitudes of currents, traces were low-pass filtered at 30 Hz.

**Optogenetic stimulation**

Light from a mercury lamp (Illuminator HXP 120 V, Zeiss) passed a GFP filter (440–490 nm band-pass) for photoactivation of cap cells co-expressing RhoGEF2-CRY2::mCherry and CIBN::pmGFP. Because chordotonal neurons are sensitive to temperature change, light pulses (400 ms, 10 $mW/mm^2$) were applied with intermittent 10 s breaks for recordings of neuronal activities, in order to avoid heat accumulation in the embedding saline solution.

**Immunostaining and microscopy of chordotonal organs**

Third instar larvae were dissected in ice-cold $Ca2^+$-free hemolymph-like saline solution (Zhang et al., 2015), and then fixed in 4% paraformaldehyde (PFA) for 30 min at room temperature, rinsed several times in 0.3% PBT (PBS with 0.3% Triton X-100, Sigma-Aldrich). To label filamentous actin, phalloidin conjugated with Atto 647N (1:400, Atto-Tec) was added to the samples and incubated overnight at 4°C. After the samples were rinsed for 4x15 min in 0.3% PBT and mounted on glass slides using Polyvinyl alcohol mounting medium with DABCO (Sigma-Aldrich). To label GFP tagged myosin, rabbit anti-GFP conjugated with Alexa Fluor 488 (Thermo Fisher Scientific) was diluted (1:500) in blocking solution and the samples were incubated overnight at 4°C. Confocal images were acquired from lch5 organs in the abdominal segments with a Leica TCS SP8 confocal microscope (Leica, Germany).

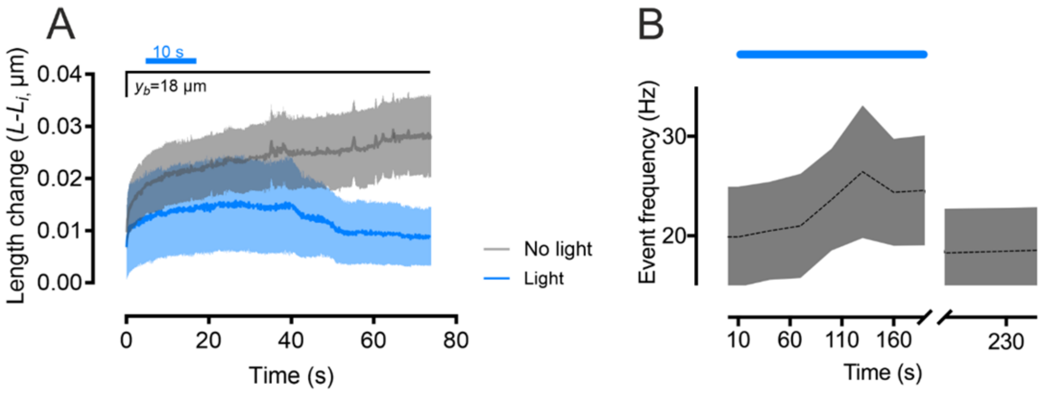


**Fig. S1. Change in relative length and firing of lch5 organ imposed by optogenetic activation of myosin. (A)** Relative length change of the organ following optogenetic activation of myosin in cap cells. Traces show the length changes of lch5 over time with (dark blue/light blue, average ± SEM, N = 5 larvae) and without (dark grey/light grey, average ± SEM, N = 3 larvae) light-induced optogenetic myosin-II activation. **(B)** Firing frequencies of lch5 neurons increase during the optogenetic stimulation that causes cap cell contractility (N = 7 larvae). Photostimulation pulse (400 ms) every 10 s. Data are presented as mean ± SEM.

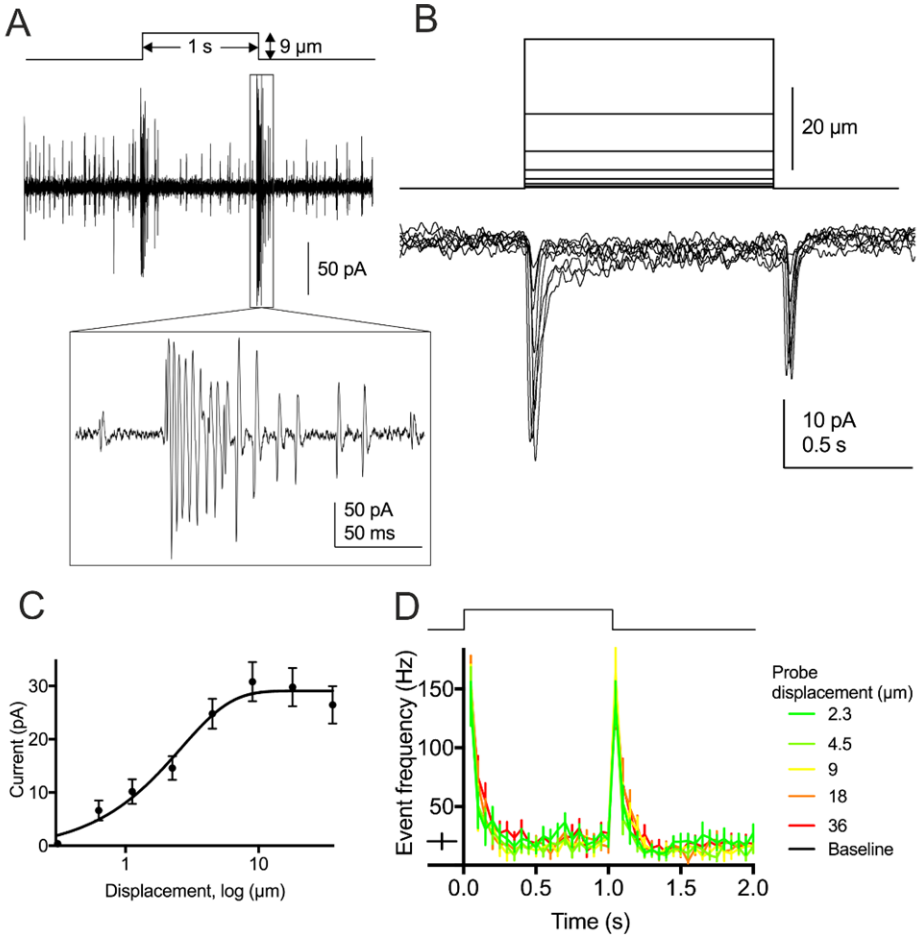


**Fig. S2. Mechano-electrical transduction of lch5 neurons. (A)** Representative recordings from an lch5 axon bundle evoked by a single-step protocol. Boxed region illustrates a burst of individual action currents. **(B)** Receptor currents from lch5 neurons during 8 step displacements from 0.31 to 36 µm (upper traces) in the presence of TTX (average of 10 sweeps). **(C)** Plot of receptor currents versus stimulus size (error bars: ± SEM, N = 10). The line is a fit with a Boltzmann sigmoidal function. A plateau is reached at displacements of ~10 µm, with maximum sensitivity occurring from 0.31 to 9 µm. **(D)** Action current frequencies (50 ms bins) for all step sizes (color coded) show robust rapidly adapting and symmetrical responses.

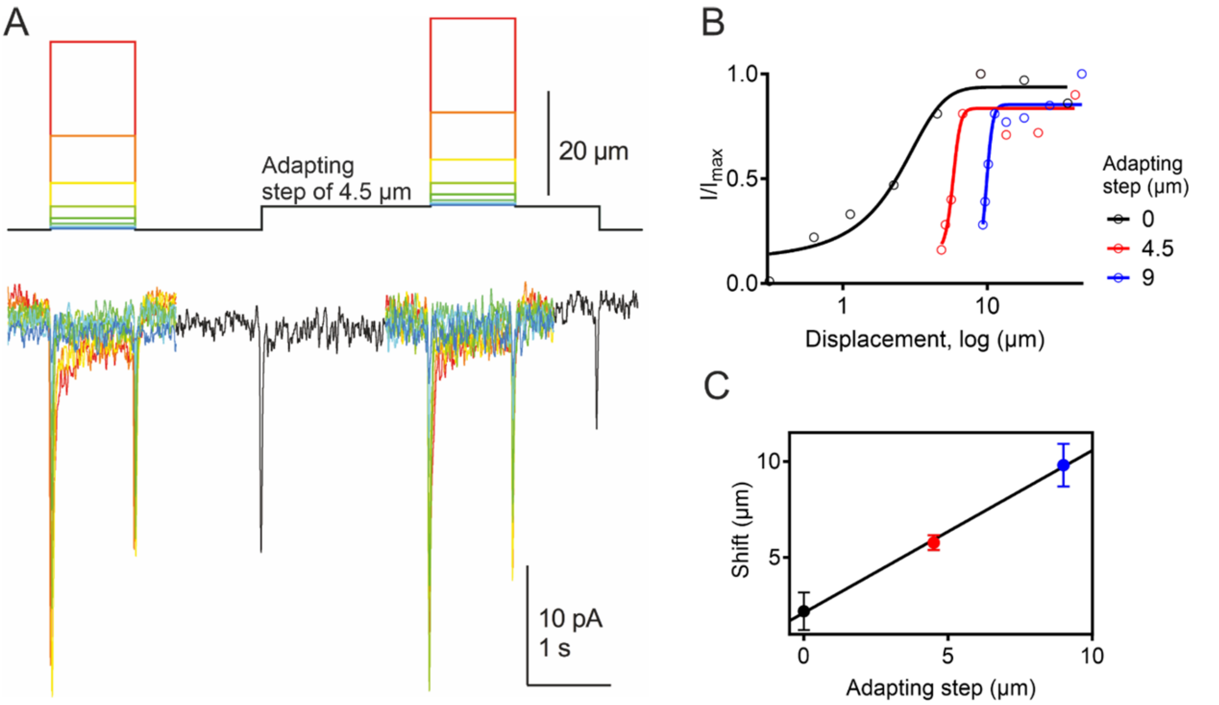


**Fig. S3. Adaptive shifts of lch5 mechanosensory transduction currents. (A)** A series of test stimuli were given before and during a 4.5 µm adapting step (upper traces) while receptor currents were recorded (lower traces). **(B)** The black and colored (red: 4.5 µm and blue: 9 µm adapting step) symbols show the current-displacement plots generated before and during adapting steps, respectively. Solid lines are single Boltzmann sigmoidal fits to the data. **(C)** The shift of each *I*(*X*) curve was plotted versus its adapting step. The linear slope (0.84) reflects the relative extent of the adaptive response. Color code as in (B).

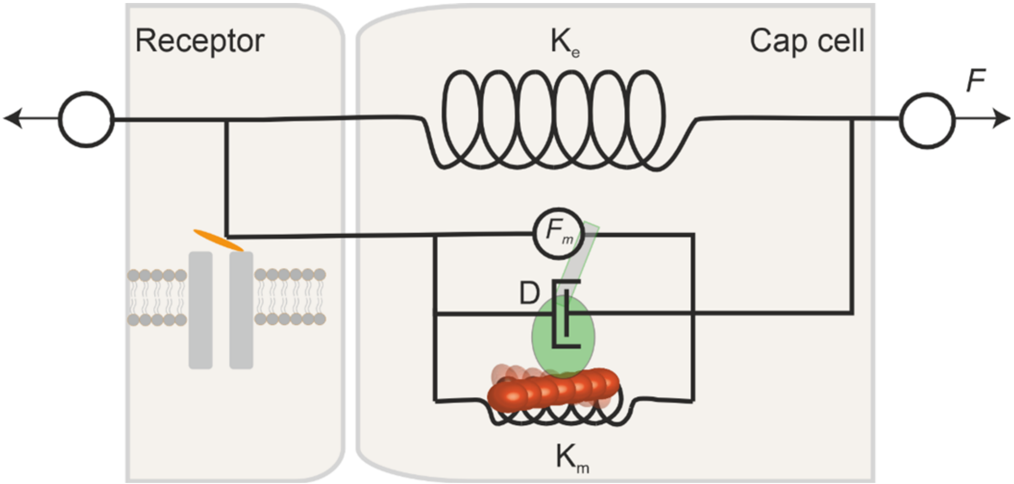


**Fig. S4. Hypothesized model for lch5 mechanosensation and adaptation involving myosin**. Tension (*F*) is an external force balanced along the stretched organ. Most of the elastic strain is localized in the cap cells, where the intrinsic stiffness is set by mechanical elements, including an extracellular matrix spring ($K_e$) and an actomyosin fiber spring (gating spring, $K_m$). A lumped damping coefficient is identified by a dashpot (D). Myosin motors of the gating spring generate force ($F_m$) that acts in parallel to external forces to exert force on the transducer channels in the sensory cilium.

**References:**


41. A. Kishino, T. Yanagida, Force measurements by micromanipulation of a single actin filament by glass needles. *Nature* **334**, 74-76 (1988).
42. M. Backholm, W. S. Ryu, K. Dalnoki-Veress, Viscoelastic properties of the nematode Caenorhabditis elegans, a self-similar, shear-thinning worm. *Proc. Natl. Acad. Sci. U.S.A.* **110**, 4528-4533 (2013).
43. M. Backholm, O. Bäumchen, Micropipette force sensors for in vivo force measurements on single cells and multicellular microorganisms. *Nat. Protoc.* **14**, 594-615 (2019).
44. C. T. Kreis, M. Le Blay, C. Linne, M. M. Makowski, O. Bäumchen, Adhesion of *Chlamydomonas* microalgae to surfaces is switchable by light. *Nat. Phys.* **14**, 45-49 (2018).
